# Inferring active mutational processes in cancer using single cell sequencing and evolutionary constraints

**DOI:** 10.1101/2025.02.24.639589

**Authors:** Gryte Satas, Matthew A. Myers, Andrew McPherson, Sohrab P. Shah

## Abstract

Ongoing mutagenesis in cancer drives genetic diversity throughout the natural history of cancers. As the activities of mutational processes are dynamic throughout evolution, distinguishing the mutational signatures of ‘active’ and ‘historical’ processes has important implications for studying how tumors evolve. This can aid in understanding mutagenic states at the time of presentation, and in associating active mutational process with therapeutic resistance. As bulk sequencing primarily captures historical mutational processes, we studied whether ultra-low-coverage single-cell whole-genome sequencing (scWGS), which measures the distribution of mutations across hundreds or thousands of individual cells, could enable the distinction between historical and active mutational processes. While technical challenges and data sparsity have limited mutation analysis in scWGS, we show that these data contain valuable information about dynamic mutational processes. To robustly interpret single nucleotide variants (SNVs) in scWGS, we introduce ArtiCull, a method to identify and remove SNV artifacts by leveraging evolutionary constraints, enabling reliable detection of mutations for signature analysis. Applying this approach to scWGS data from pancreatic ductal adenocarcinoma (PDAC), triple-negative breast cancer (TNBC), and high-grade serous ovarian cancer (HGSOC), we uncover temporal and spatial patterns in mutational processes. In PDAC, we observe a temporal increase in mismatch repair deficiency (MMRd). In cisplatin-treated TNBC patient-derived xenografts, we identify therapy-induced mutagenesis and inactivation of APOBEC3 activity. In HGSOC, we show distinct patterns of APOBEC3 mutagenesis, including late tumor-wide activation in one case and clade-specific enrichment in another. Additionally, we detect a clone-specific increase in SBS17 activity, in a clone previously linked to recurrence. Our findings establish ultra-low-coverage scWGS as a powerful approach for studying active mutational processes that may influence ongoing clonal evolution and therapeutic resistance.

## 1 Introduction

Ongoing mutagenesis in cancer is fundamental to generating genetic diversity and conferring phenotypic properties to tumor tissues. Specific endogenous and exogenous mutational processes damage or alter the genome in distinct ways, leaving characteristic *mutational signatures*^1^ in the form of specific patterns of mutation. Computational tools to identify these signatures from mutations detected in DNA sequencing data can therefore infer the underlying mutagenic processes that were active in the development of a tumor^2^. These analyses have been key for characterizing tumor evolution, understanding clonal dynamics, identifying mutational processes that confer drug resistance, and mutational signatures that serve as biomarkers for treatment decisions^3–8^. A critical challenge is that many mutational signatures observed at the time of diagnosis reflect past mutagenic activity and may not capture the *active* processes that drive or reflect current tumor behavior at the time the tumor was sampled. Consequently, mutations from ‘active’ mutational processes will likely only be present in only a small fraction of tumor cells (i.e., located near the leaves of the phylogenetic tree). Accurately recovering these low-prevalence variants and characterizing their distribution within the tumor is challenging due to the limited sensitivity of sequencing technologies^9^.

As active mutational processes reflect the tumor phenotype at time of sampling, distinguishing active from inactive mutational processes could indicate mechanisms of drug resistance and assessing their relevance as therapeutic targets. For example, in estrogen receptor-positive (ER+) breast cancer, resistance to CDK4/6 inhibitors has been linked to active APOBEC3 mutagenesis, which is associated with shorter progression-free survival and therapy resistance^10^. Similarly, in EGFR-mutant lung cancer, targeted therapies such as tyrosine kinase inhibitors (TKIs) have been shown to induce active APOBEC3A activity, leading to sustained mutagenesis and genomic instability^11^. In advanced bladder cancer, chemotherapy-induced bursts of subclonal mutations, and complex structural variants, including extrachromosomal DNA (ecDNA) amplifications, play a critical role in the evolution of resistance^12^.

One major challenge to reasoning about active mutational processes is tumor heterogeneity due to clonal evolution^13^. Tumors are composed of multiple distinct cell populations, or *clones*, that harbor shared somatic mutations from their ancestral lineage, but additional shared mutations within the population that are mutually exclusive with other clones. As tumors evolve under selective pressures from the microenvironment, immune system, and treatment, distinct clonal populations may activate endogenous processes which can lead to the emergence of increasingly aggressive or drug-resistant clones^14–18^. Knowing which clone (or clones) harbor which mutations is critical to understanding active mutational processes. Single-cell DNA sequencing (scDNA-seq) has great potential to uniquely to address this issue, enabling the measurement of genetic variation and evolutionary processes within individual cells of a tumor^19^. Typical approaches center on targeted and exome scDNA-seq experiments to study SNVs as they provide provide high per-cell read depth, resulting in precise SNV identification^20, 21^. However, these approaches sacrifice coverage breadth: they capture only a small portion of the genome (≤1%), which greatly limits the number of observable variants. This limitation hinders the scope of SNV-based analyses, including studies of active mutational processes, mutation rates, and evolutionary patterns.

An intriguing alternative for SNV-based analyses is single-cell whole-genome sequencing (scWGS) at ultra-low coverage^22–26^, which is typically used for the study of copy-number alterations. Although the low coverage (*<*0.25×) precludes confident SNV calling in individual cells, the whole-genome breadth enables identification of large numbers of SNVs across the cell population. By aggregating data across cells into clones, ultra-low-coverage scWGS provides a potential means to analyze clonal tumor populations and their evolutionary dynamics, leveraging both SNV and CNA data. This approach is particularly valuable for studying active mutational processes, as examining mutational profiles at the clonal level can reveal the late emergence of mutational signatures. Such signatures, which may appear at low frequencies, are often obscured by signal averaging inherent in bulk sequencing approaches.

SNV calling in scWGS data remains challenging due to the high rates of technical artifacts^20^. Errors introduced during sample preservation, library preparation, amplification, and sequencing can obscure true biological signals. While single-cell-specific methods such as ProSolo^27^, Monovar^28^, and SCcaller^29^, these methods depend on the high per-cell coverage of targeted and exome sequencing data. Comparable tools do not exist for ultra-low coverage scWGS data, where most cells have too few reads to allow confident SNV calls at the individual cell level. As a result, single-cell data is typically pooled into pseudo-bulk samples which can be analyzed by standard bulk somatic variant callers^30^. While pseudo-bulk approaches provide a practical solution for initial variant calling, the distinct error profiles of single-cell data require additional refinement to reduce false positives and improve the overall accuracy of the variant set. This refinement process is crucial for enabling more accurate analyses of mutational signatures and tumor evolution. Motivated by the biological problem of inferring active mutational processes, in this manuscript, we address the problem of variant call refinement for low-coverage scWGS data.

We introduce ArtiCull (<u>Arti</u>fact <u>Cull</u>er), a novel variant call refinement algorithm specifically designed for ultra-low-coverage single-cell DNA sequencing data. ArtiCull imposes evolutionary constraints that are valid for any phylogeny under the evolutionary model and obviates the expense and complexity of simultaneous phylogeny inference. Central to ArtiCull’s approach is the assumption that all true variants are generated by an underlying evolutionary process reflected in a phylogeny, whereas artifacts occur independently of this process. We show how this enables robust inference of mutations, including low-prevalence and clone specific SNVs, and active mutational processes. Through analysis of scWGS from three human cancer types, we reveal biologically relevant shifts in mutational processes: a temporal increase in MMRd in PDAC, chemotherapy-induced mutagenesis and APOBEC3 inactivation in TNBC, distinct patterns of APOBEC3 activity in HG-SOC, including late tumor-wide activation, clade-specific activity, and a clone-specific increase in SBS17 linked to recurrence.

## 2 Results

### 2.1 ArtiCull: improving SNV detection in ultra-low-coverage scWGS

ArtiCull has two key components: (1) a method to identify a subset of putatively artifactual SNVs from scWGS data based on evolutionary principles (Fig. 1A, B); (2) a supervised model trained on variants labeled by (1) (Fig. 1C, D). We briefly describe these components below and elaborate in Section 4.1-4.4.

**Figure 1:**
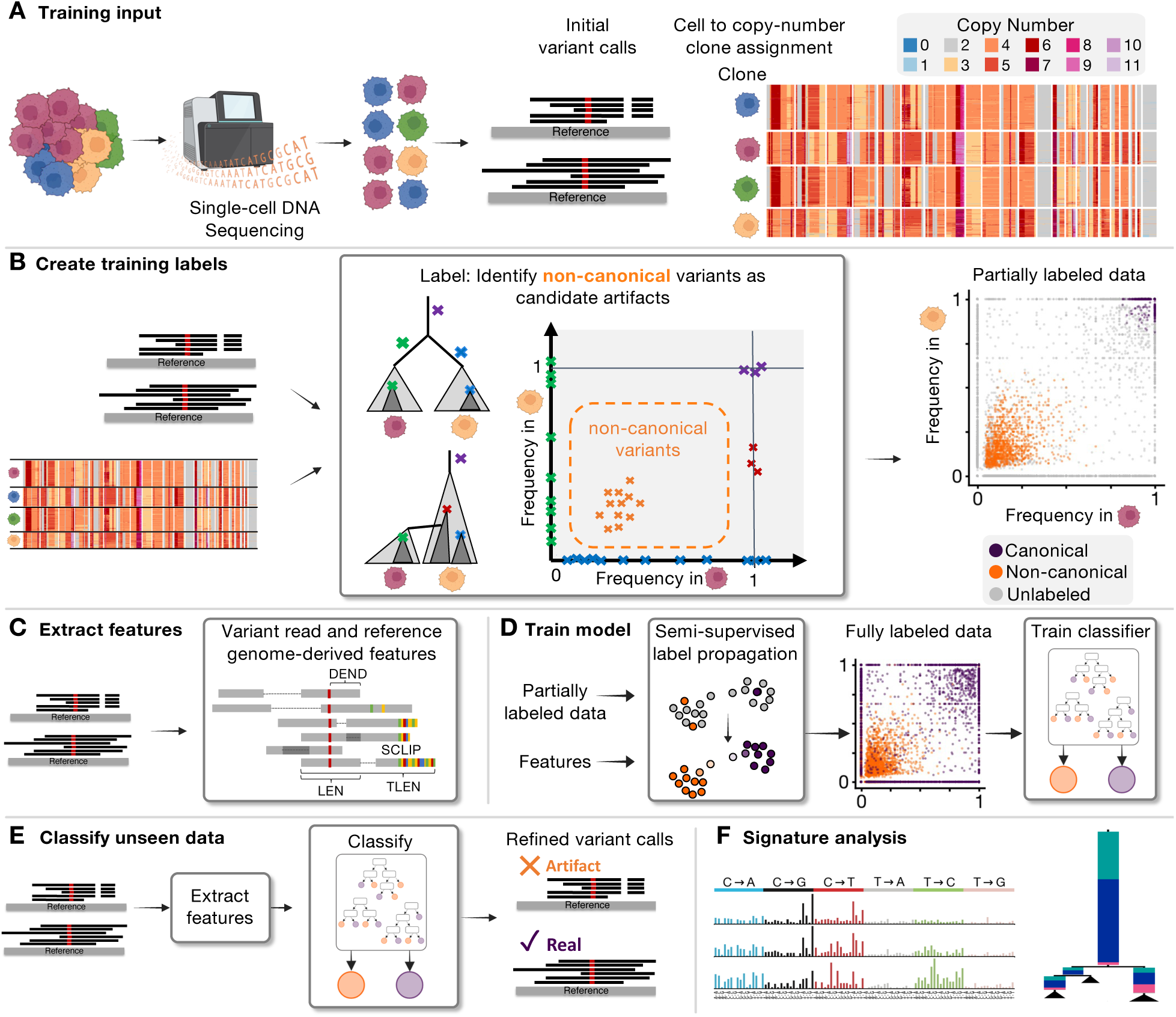
Overview of the ArtiCull Method and Training Process. ArtiCull is a method designed to enhance the accuracy and reliability of variant calling in single-cell whole-genome sequencing by identifying and filtering candidate artifacts. **(A)** *Data Setup and Input*. Training the ArtiCull model begins with a set of initial variant calls derived from single-cell sequencing data and labels that assign cells to copy-number clones based on single-cell copy-number profiles. **(B)** *Training Label Generation*. Using the input variant calls and clone labels, ArtiCull identifies candidate artifacts (non-canonical variants) alongside clonal variants that are candidate real mutations yielding a partially-labeled dataset. **(C)** *Feature Extraction*. Features are extracted directly from sequencing data. These features include read-level metrics and genome-derived characteristics for each variant call. **(D)** *Model Training*. The partially labeled data (Panel B) and the features (Panel C) are used to train a classification model. Semi-supervised label propagation is first applied to generate fully labeled data, which is then used to train a gradient-boosted tree classifier. **(E)** *Application to Unseen Data*. The trained classifier is applied to new variant call datasets. Features are extracted from these datasets, and the classifier refines the variant call set by identifying and removing candidate artifacts. Note that this process does not take copy-number information as input. **(F)** The refined variant call set can be used for downstream analyses, including mutation signature analysis.

As previously mentioned, we impose the assumption that all true variants are generated by an underlying evolutionary process reflected in a phylogeny, whereas artifacts occur independently of this process. This assumption allows us to identify non-canonical variants—those that could not have been generated by or do not respect the evolutionary process. One approach for identifying such variants, which has been used by earlier methods^31, 32^, is to construct a full phylogeny and flag variants that do not fit this phylogeny. However, phylogeny construction is computationally expensive and error-prone, particularly in sparse data, limiting both accuracy and scalability. Below, we define an evolutionary constraint that holds for *any* phylogeny consistent with an evolutionary model and can be efficiently evaluated for each variant.

To identify non-canonical variants, we incorporate orthogonal copy-number data from the same cells to establish *copy-number clones* (sets of cells with identical copy-number profiles), which hereafter we refer to as clones for simplicity. The *clone-specific cell fraction (CF) c_A_* of an SNV is the proportion of cells in clone *A* that contain a mutation. An SNV is *clonal* with respect to clone A if *c_A_* = 1, *subclonal* if 0 *< c_A_ <* 1, and *absent* if *c_A_* = 0.

#### Definition 1

A variant is canonical *if there exists at most one clone A in which* 0 *< c_A_ <* 1*, and* non-canonical *otherwise*.

In Section 4.1, we prove that under common assumptions about somatic evolution, non-canonical variants cannot derive from any valid phylogeny. In noisy sequencing data where CFs are not directly observed, ArtiCull identifies confidently canonical and non-canonical variants using a hypothesis test approach and a probabilistic sequencing model (see Section 4.2). However, this approach alone is insufficient for identifying all artifacts; some artifacts—particularly those at low frequencies—may conform to a phylogeny by chance, making them incidentally canonical. To address this limitation, we first classify variants as either confidently non-canonical, canonical, or uncertain (Fig. 1B). Using variant read and reference genome-derived features (Fig. 1C), these labeled variants are then used to train a model to distinguish artifacts from true SNVs (Fig. 1D). Once trained, this model can be applied to unseen data from new single-cell sequencing samples (Fig. 1E). Copy-number data is only used generate the initial training labels; neither the model training nor its application on new data observes copy-number information.

### 2.2 Characterizing non-canonical variants and evaluating classifier performance

We first evaluated the assumption that non-canonical variants are likely artifacts. Variants from seven breast and ovarian DLP+ samples were labeled as canonical, non-canonical, or unlabeled (as described in Section 4.4). Non-canonical variants displayed characteristics typical of artifacts, including read-level anomalies and genome distribution patterns^33^. ArtiCull extracts a set of fifteen read and genome-level features summarized in Table 1, Fig. 2A. Non-canonical variants showed significant differences from canonical variants across all fifteen features (Mann-Whitney U test, *p* ≤ 0.003; Fig. 2A). For example, non-canonical variants had lower mapping qualities (MAPQ: medians 25.8 vs. 60, *p <* 0.001), lower variance in start and end positions (SSTD: 14.1 vs. 40.6, *p <* 0.001; ESTD: 13.8 v 40.5, *p <* 0.001), shorter template lengths (TLEN: 127 vs. 271, *p <* 0.001), and more base mismatches (MM: 2.2 vs. 1.5, *p <* 0.001). These features effectively separate canonical from non-canonical variants (Fig. 2B; See Fig. S4 for per-sample and per-feature distributions).

**Figure 2:**
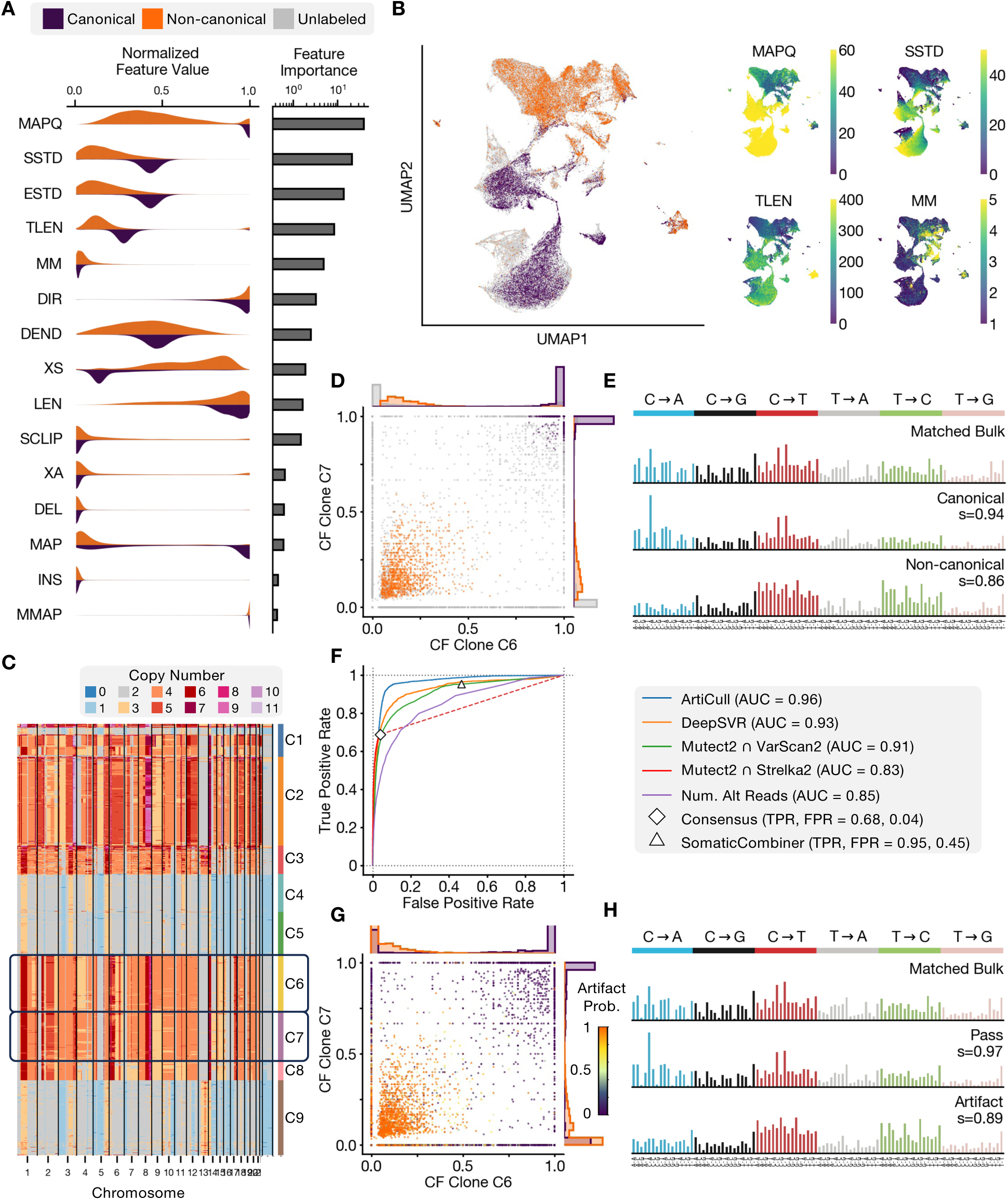
Benchmarking and evaluation of ArtiCull frequency-based labeling and feature-based classification. **(A)** Variant calls from seven ovarian and breast cancer samples were used to train an ArtiCull model. Feature distribution for labeled canonical (purple) and non-canonical (orange) variants from the training dataset. Feature definitions are provided in Table 1, and feature importance for the trained model is shown alongside the distribution. **(B)** UMAP visualization of training variant calls based on extracted features. **(left)** Points are colored based on their label as canonical (orange), non-canonical (purple) or unlabeled (gray). **(right)** UMAPs colored for four of the highest importance features: MAPQ (mapping quality), SSTD (start position standard deviation), TLEN (template length), and MM (number of mismatches). **(C)** Copy number visualization for cells in OV2295. Copy number and cluster labels are derived from SIGNALS. **(D)** Pairwise clone-specific cell fraction (CF) comparison for clones C6 and C7 from OV2295. The hypothesis test was used to label the points based on the CFs (colored same as panel B). Histograms along the axes show per-cluster CF distribution with respect to these labels. **(E)** Mutation signature distributions for matched bulk, canonical, and non-canonical variants. Cosine similarity (*s*) indicates the similarity between a distribution and the matched bulk distribution. **(F)** Model performance was evaluated on a held-out ovarian cancer sample (OV-022). Variant calls from a site-matched bulk tumor sample was used as ground truth. Receiver-Operator Curves (ROC) are shown for ArtiCull results as well as a set of alternative variant refinement methods. Neither the consensus approach (an intersection of Mutect2, Strelka2 and VarScan2) or SomaticCombiner (using results from Mutect2, Strelka2 and VarScan2) provide a per-variant score, and thus are represented as individual points. **(G)** The pre-trained ArtiCull model was applied to OV2295. The resulting artifact probabilities are indicated on the CF distribution by color. Histograms along the axes indicate mutations identified as artifacts (artifact probability *>* 0.5, in orange) and mutations that pass (prob. ≤ 0.5 in purple). **(H)** Mutation signature distributions for bulk, pass, and artifact variants, with cosine similarity as described in panel F.

**Table 1:**
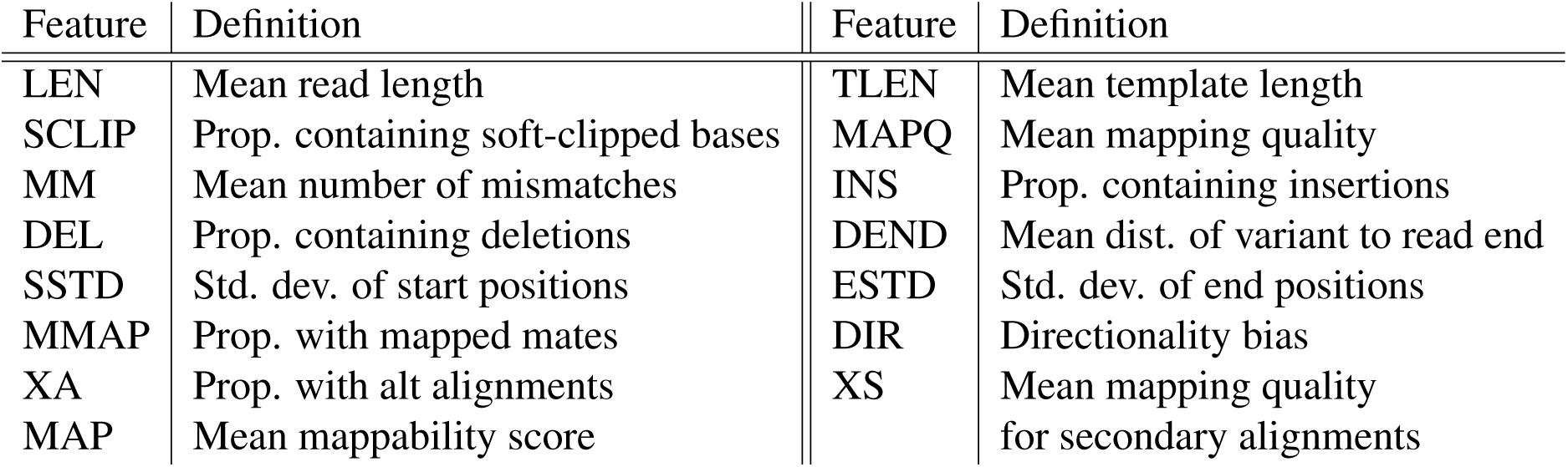
Model features. Brief descriptions of model features. All features except MAP are computed over the set of variant reads aligned at the variant position. Formal definitions are provided in Section S1.

| Feature | Definition | Feature | Definition |
| --- | --- | --- | --- |
| LEN | Mean read length | TLEN | Mean template length |
| SCLIP | Prop. containing soft-clipped bases | MAPQ | Mean mapping quality |
| MM | Mean number of mismatches | INS | Prop. containing insertions |
| DEL | Prop. containing deletions | DEND | Mean dist. of variant to read end |
| SSTD | Std. dev. of start positions | ESTD | Std. dev. of end positions |
| MMAP | Prop. with mapped mates | DIR | Directionality bias |
| XA | Prop. with alt alignments | XS | Mean mapping quality for secondary alignments |
| MAP | Mean mappability score |  |  |

We next compared the detection rates of canonical and non-canonical variants in matched bulk sequencing data from two ovarian cancer datasets (OV2295: Fig. 2C-E; OV-022: Fig. S1). For the two datasets, only 1.7% and 1.1% of non-canonical variants were called in the bulk sequencing data, respectively, compared to 91% and 95% of canonical variants. For unlabeled variants, 12% and 13% of unlabeled variants were detected in the bulk data, despite unlabeled variants having lower allele frequencies and numbers of variant read counts than non-canonical variants on average. We further analyzed the 96-channel trinucleotide mutational profiles of bulk sequencing data, finding that non-canonical variants exhibited lower cosine similarity to the bulk mutational profile than canonical variants (OV2295: Fig. 2G; OV-022: Fig. S1). Taken together, these three lines of evidence support that non-canonical variants identified by ArtiCull are predominantly artifacts rather than true mutations.

Variant labels from seven breast and ovarian cancer DLP+ sequencing samples were used to train a classifier (as described in Section 4.4). To evaluate the performance of the trained classifier, we computed its performance using a held-out sample with site-matched bulk sequencing (OV-022), and compared against other variant call refinement approaches (Fig. 2F). These approaches included: (1) taking the intersection of the initial variant call set (using Mutect2^34^) with other variant callers, Varscan2^35^ and Strelka2^36^. Variants were ranked variants using the reported somatic p-value by Varscan2 and the QSS (Quality Score for SNV) for Strelka2; (2) SomaticCombiner^37^, an ensemble approach for variant refinement using Mutect2, Varscan2, and Strelka2 as the input call set; (3) DeepSVR^38^, a feature-based classifier approach trained on a large cohort of bulk sequencing data; and (4) ranking variants based on the number of observed variant reads. ArtiCull displayed the best performance, achieving an AUC of 0.96 (Fig. 2F). DeepSVR, which uses a similar feature-based approach as ArtiCull, likewise performed well (AUC = 0.93). The difference in performance may reflect ArtiCull being better tuned to the feature distribution for these samples due to being trained on scWGS. Notably, DeepSVR was trained on manually annotated data which required nearly 600 hours of expert labor^38^. ArtiCull in contrast achieves this performance with no manual labeling. The intersection of Mutect2 and VarScan2 also performed competitively (AUC = 0.91), while other methods had lower performance.

### 2.3 Mutational dynamics in pancreatic ductal adenocarcinoma with mismatch repair deficiency

To demonstrate inference of active mutational processes in scWGS, we analyzed DLP+^23^ scWGS of patient-derived organoids from a mismatch repair deficient (MMRd) pancreatic ductal adenocarcinoma (PDAC) sample. We used SBMClone^39^, a method specifically designed for bi-clustering cells and mutations in highly sparse variant call data such as scWGS, to analyze ArtiCull-filtered variant calls (Section 4.5). These samples show substantial intratumor heterogeneity with nine distinct cell clusters defined by thirteen SNV clusters (Fig. 3A). The robustness of the SNV-based clustering is supported by its concordance with copy-number profiles in the same cells (Fig. 3B).

**Figure 3:**
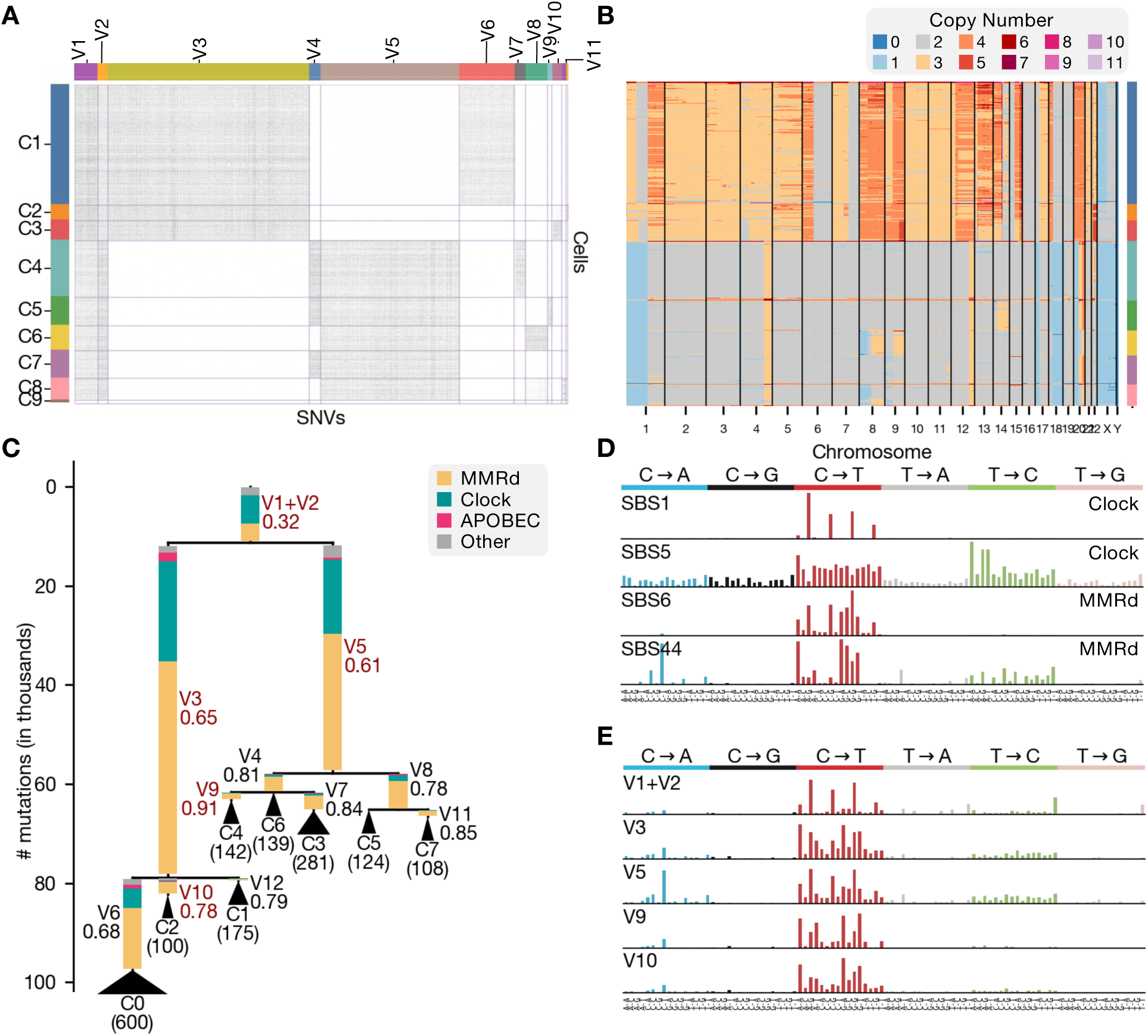
Mutation signature analysis of mismatch repair deficient pancreatic adenocarcinoma. **(A)** SBMClone was used to cluster cells and mutations and identified nine clusters of cells (rows), defined by thirteen clusters of SNVs (columns). **(B)** Heatmap of the copy number profiles for the cells, sorted in the same order as cells in panel A. **(C)** Phylogeny relating the clusters of cells from the SBMClone clustering. MuSiCal was used to refit the sets of mutations in each SNV cluster from panel A, with resulting signatures grouped as Clock-like (cyan), APOBEC (pink), mismatch repair deficiency (MMRd; yellow), and Other (gray). Each edge corresponds to one SNV cluster, with values under the SNV labels indicating the proportion of mutations attributed to MMRd-like signatures. Red labels correspond to select SNV clusters shown in panel D. Each leaf is labeled by a cell cluster from panel A, with the number in parentheses indicating the number of cells in that cluster. **(D)** COSMIC v3.4 mutation signatures. Clock-like signatures SBS1 and SBS5 are shown, along with MMRd signatures SBS6 and SBS44, the two most prevalent MMRd signatures in these samples. **(E)** Mutation signature distributions for select SNV clusters. V1+V2 correspond to truncal mutations. V3 and V5 correspond to intermediate branches. V9 and V10 correspond to branches near the leaves.

From the SBMClone clustering results, we constructed a phylogeny using perfect phylogeny principles^40^, which revealed a bifurcated structure (Fig. 3B). When integrated with copy number data, we found these two major clades corresponded to distinct ploidy states: one clade with a predominantly diploid state, while the other had predominantly triploid and tetraploid chromosomes likely resulting from an earlier whole-genome duplication. Mutational signature decomposition using MuSiCal^41^ with COSMIC single-base substitution signatures (SBS) v3.4 (Section 4.6) revealed distinct temporal patterns. We classified the identified signatures as Clock-like (SBS1, 5), MMRd-associated (SBS6, 14, 15, 21, 26, 44), APOBEC3-associated (SBS2, 13) and Other. Our analysis revealed increasing MMRd activity across both clades (Fig. 3B-D): truncal mutations had relatively lower MMRd activity (32%), which progressively increased in intermediate branches (65% and 61%) and reached a peak in the terminal branches (68%-91%). The WGD clade showed somewhat higher overall mutational burden, consistent with increased mutational opportunity from higher DNA content, but followed the same pattern of intensifying MMRd activity. This analysis demonstrates how single-cell sequencing can reveal the temporal dynamics of mutational processes during tumor evolution.

### 2.4 Emergence of cisplatin-associated mutations in triple-negative breast cancer

Since mutation signature decomposition typically lacks a definitive ground truth, we aimed to assess whether ArtiCull could identify mutational signatures consistent with known mutagenic processes, such as those induced by platinum-based chemotherapy. Platinum-based chemotherapy such as cisplatin is known to induce characteristic mutational signatures, particularly SBS31 and 35^42^. To evaluate the ability of ArtiCull to capture temporally active mutational signatures, we applied ArtiCull to variant calls from sequential passages of a triple-negative breast cancer (TNBC) patient-derived xenograft (PDX) experiment^43^ SA609 (Fig. 4A). The first sample, X3-Untreated, corresponds to a treatment-naive tumor. In the subsequent passages (X4-Rx to X7-Rx), mice were treated with sublethal dosing of cisplatin between each passage. A parallel control sequence of passages was completed with no treatment between passages (X4-U to X7-U). Samples X4-Rx and X5-U were excluded from analysis due to low high-quality cell counts (*<*50 cells). Given the cumulative exposure to chemotherapy, we expected to observe progressive accumulation of cisplatin-associated mutations in the passages, and no such accumulation in the untreated samples.

**Figure 4:**
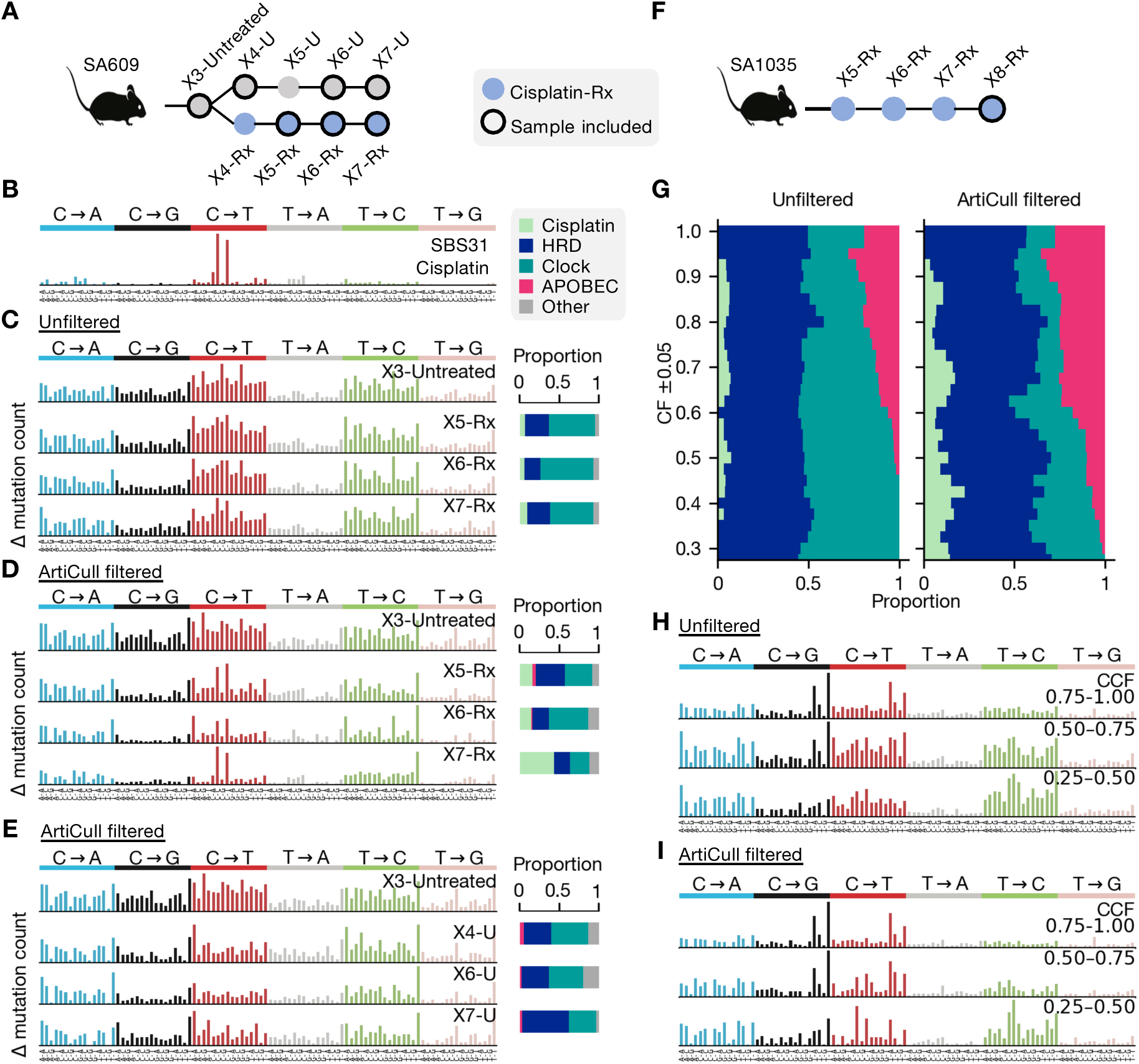
Mutation Signature Analysis Across Sequential Passages of TNBC PDX Experiments. **(A)** The SA609 TNBC experiment consists of sequential passages of a TNBC patient-derived xenograft (PDX). Samples originate from X3-Untreated, after which the experiment branches into two arms: a treated arm (X4-Rx, X5-Rx, X6-Rx, X7-Rx), where cisplatin was administered between passages, and an untreated arm (X4-U, X5-U, X6-U, X7-U) without treatment. Samples X4-Rx and X5-U were excluded from analyses due to low cell counts. **(B)** Cisplatin-associated signature SBS31. **(C-E)** Mutation profiles are shown for the untreated baseline sample (X3-Untreated) and subsequent passages. Each profile represents mutations accumulated since the previous time point. Panels **(C)** and **(D)** show profiles for the cisplatin-treated arm (X5-Rx, X6-Rx, X7-Rx), using unfiltered and ArtiCull-filtered variant call sets, respectively. Panel (E) shows ArtiCull-filtered profiles for the untreated arm (X4-U, X6-U, X7-U). **(F)** The SA1035 TNBC PDX comprises sequential passages of a TNBC patient-derived xenograft (PDX). Cisplatin was administered between passages. **(G-H)** Mutation profiles for X8-Rx unfiltered **(G)** and ArtiCull-filtered **(H)** data, categorized by clone-specific cell fraction (CF) bins (0.25 0.50, 0.50 0.75, 0.75 1.00). **(I)** Moving window signature decomposition for X8-Rx displaying the distribution of mutational signatures (Clock-like, HRD, APOBEC3, and Cisplatin-associated) across different CF ranges, for unfiltered (left) and ArtiCull filtered (right) variant call sets.

Using X3-Untreated as a baseline, we analyzed the mutations that were newly acquired in each sample relative to its immediate predecessor. MuSiCal was used to perform signature decomposition and signatures were classified as Clock-like (SBS1, 5), homologous recombination deficiency (HRD) associated (SBS3, 8), APOBEC (SBS2, 13), cisplatin (SBS31, 35), and Other. SBS35 was not detected in any samples, and thus all cisplatin exposure was attributed to SBS31 (Fig. 4B).

In the unfiltered data (Fig. 4C), the cisplatin signature was not evident and the signature decomposition estimated activity in low amounts (6%, 5% and 9% respectively for X5-Rx, X6-Rx, and X7-Rx). After applying ArtiCull (Fig. 4D), the cisplatin-associated mutations are more evident, and the decomposition estimated activity at 15%, 14% and 43%. As a control, we also examined a parallel arm of passages that were not subject to cisplatin treatment (Fig. 4E). In these samples, no cisplatin activity was inferred in either the filtered or unfiltered data.

We next aimed to demonstrate how ArtiCull impacts the analysis of low-frequency, late-occurring variants within samples, which are crucial for understanding mutational processes active during late-stage tumor evolution. These variants are often obscured by artifacts, making it challenging to detect shifts in mutational process activity over time. For this analysis, we examined a second TNBC PDX transplant experiment, SA1035, from Salehi et al.^43^, (Fig. 4F). We focused on a sample from passage 8 (X8-Rx) which had received four rounds of cisplatin exposure. To evaluate the sensitivity to detect shifts in signatures and late-emerging mutation signatures, we used a moving-window signature decomposition over CF values to bin mutations (i.e., mutations with CF between 0.9 and 1.0, then CF between 0.875 and 0.975, etc.; Fig. 4G). Sets with higher CF likely contain earlier-acquired mutations present in more cells, whereas sets with CF indicate more recent mutagenesis. In both the filtered and unfiltered data, this sample exhibited strong evidence of both HRD and APOBEC3-associated signatures, where APOBEC3 decrease at lower CF (Fig. 4G). However, the filtered and unfiltered data differed with respect to cisplatin-associated signatures: the ArtiCull filtered data showed an increase in cisplatin-associated signatures as frequency decreases, whereas in the unfiltered data the cisplatin signature was either present in low amounts or completely absent. Additionally, in the unfiltered data at lower CF, we observed an increased assignment of mutations to clock-like signatures, particularly SBS5. This was likely due to the relatively flat profile of SBS5, which makes it more prone to absorbing noise. While we lack ground truth for these samples, the filtered data aligns more closely with the known treatment history of this PDX series.

### 2.5 Late activation of APOBEC3 mutagenesis in high-grade serous ovarian cancer

We analyzed 654 cells from two DLP+ scWGS samples from high-grade serous ovarian cancer patient OV-046 from McPherson et al.^44^(Fig. 5A). To evaluate temporal shifts in mutational signatures, we applied a moving-window signature decomposition over mutation CF ranges(as described in Section 2.4; Fig. 5B). In the ArtiCull-filtered data, ongoing APOBEC3 activity (Fig. 5C) was detectable and visually evident even at low CF (Fig. 5D). However, in the unfiltered data, this APOBEC3 activity was largely obscured at low frequencies, with increasing numbers of mutations being attributed to the flat clock signature SBS5.

**Figure 5:**
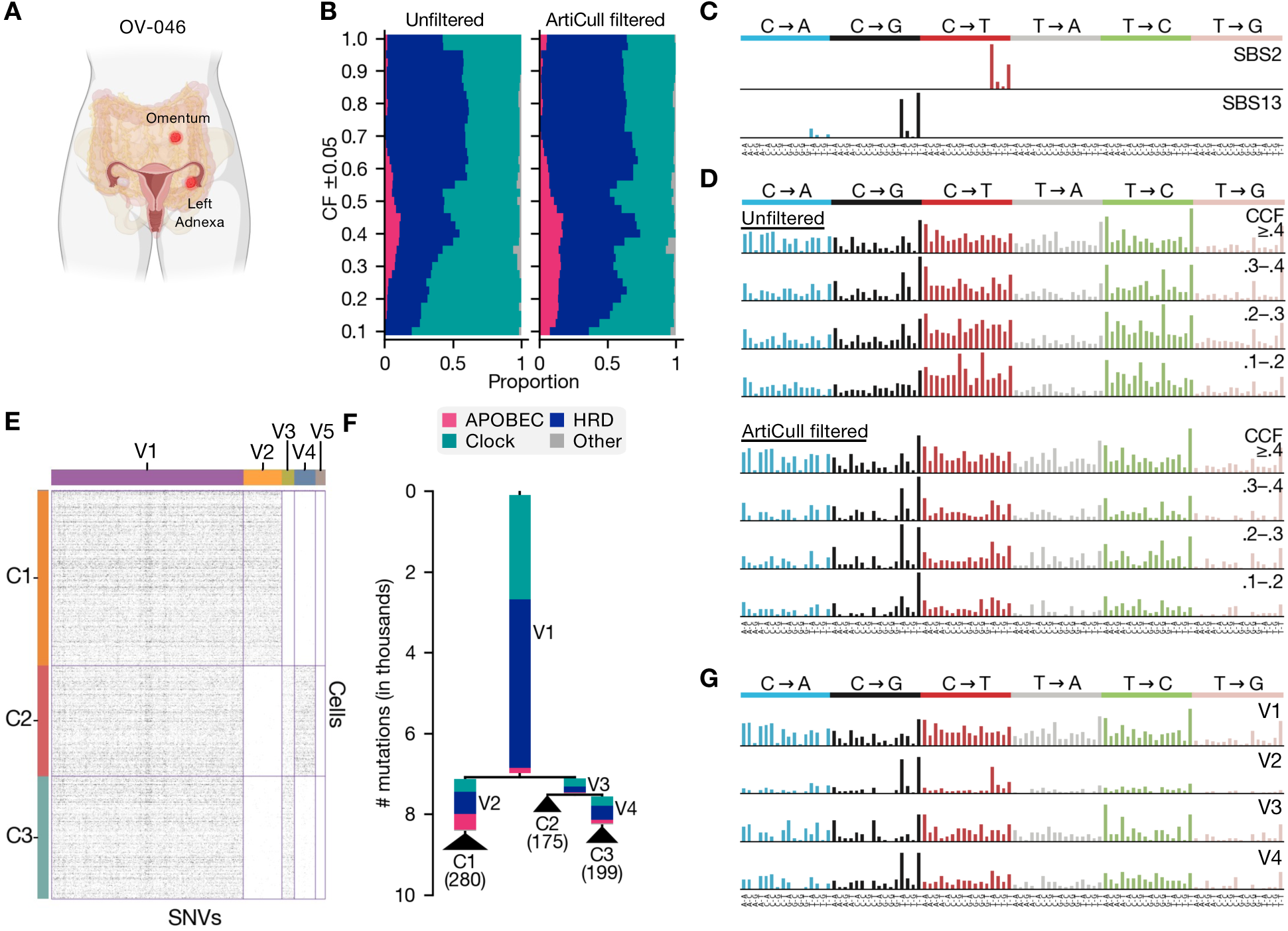
Clustering and Mutational Signature Analysis of High-Grade Serous Ovarian Cancer (HGSOC) Sample OV-046. **(A)** Overview of HGSOC OV-046, including samples from the infracolic omentum and left adnexa. **(B)** Moving window decomposition of mutational signatures across clone-specific cell fraction (CF) ranges in unfiltered (left) and ArtiCull-filtered (right) data, demonstrating shifts in mutational processes in lower-frequency variants. **(C)** APOBEC3 associated mutation signatures SBS2 and SBS13 (COSMIC mutational signatures v3.4). **(D)** Mutation profiles across different CF bins (≥0.4, 0.3 0.4, 0.2 0.3, 0.1 0.2) for the unfiltered (top) and ArtiCull-filtered (bottom) variant call sets. **(E)** SBMClone clustering results for the ArtiCull-filtered variant call sets, identifying clusters of cells based on shared SNVs. See Fig. S3 for the unfiltered variant call set. **(F)** Phylogenetic tree relating the cell clusters from the ArtiCull-filtered data. Each edge corresponds to an SBMClone SNV cluster (panel E). Edge lengths correspond to the number of mutations, and edge colors represent the mutational signatures attributed to those mutations. Leaf labels correspond to the SBMClone cell cluster labels (panel E). Numbers in parentheses indicate the number of cells in each cell cluster. **(G)** Mutation profiles of SNVs assigned to different SNV clusters in the ArtiCull-filtered data.

To evaluate how mutations are distributed with respect to the subclonal structure of this cancer, we used SBMClone^39^ to identify SNV-based clusters of cells in both the unfiltered and ArtiCull-filtered data. In the unfiltered dataset, SBMClone identifies six distinct clusters of cells (Fig. S3A). To assess the validity of these clusters, we cross-referenced the orthogonal copy-number information from these cells (Fig. S3B), and observe that the profiles do not support the proposed divisions. In contrast, the ArtiCull-filtered data (Fig. 5E) resulted in three clusters that more effectively grouped the cells based on their underlying copy-number profiles. This suggests that the erroneous clustering in the unfiltered data may have been be driven by sample-specific artifacts which were eliminated by ArtiCull. A phylogeny (Fig. 5I) was constructed relating these cell clusters (Section 4.5) and MuSiCal was used to decompose mutation signatures for each SNV cluster. Truncal mutation cluster V1, and cluster V3 (ancestral to cell clusters C1 and C2) show low levels (1.7% and 2.4%, respectively) of APOBEC3-associated signatures, SBS2 and SBS13 (Fig. 5C. In contrast, the terminal branches V2 and V4 showed a marked increase in APOBEC3 activity (13% and 29%). The near absence of APOBEC3 signatures in V3 suggests that the increase in APOBEC3 mutagenic emerged tumor-wide post-divergence of the three clones.

### 2.6 Clone-specific shifts in mutational processes in high-grade serous ovarian cancer

We analyzed 1921 cells from four DLP+ scWGS samples from high-grade serous ovarian cancer patient OV-045 from McPherson et al.^44^ (Fig. 6A). SBMClone was used to identify SNV-based clusters of cells in the ArtiCull-filtered data, revealing four cell clusters (C1–C4) and seven SNV clusters (V1–V7) (Fig. 6B). These clusters were consistent with the copy-number profiles of the constituent cells (Fig. 6C). We constructed a phylogenetic tree based on the SBMClone clusters (Fig. 6D) and used MuSiCal to decompose clone-specific mutational profiles. Signatures were categorized as HRD-associated (SBS3, 8), APOBEC3 (SBS2, 13), Clock (SBS1, 5), SBS17 (SBS17a, 17b) and Other. The signature decomposition reveals a clade-specific increase in APOBEC3 activity. Cluster V5, which is the terminal branch leading to cell cluster C1, maintained a similar level of APOBEC3 activity as truncal mutations V1 and V2 (10% vs. 11%, respectively). In contrast, APOBEC3 levels were elevated in intermediate branch V3, which precedes the divergence of clusters C2 and C4 (17%), as well as in terminal branches V4 (C4) and V6 (C3) (34% and 22%, respectively). Although the phylogeny construction based on SBMClone clustering does not group C2–4 into a distinct clade, the copy number profiles of the cells (Fig. 6C) suggest that C2–4 are more closely related to each other than to C1.

**Figure 6:**
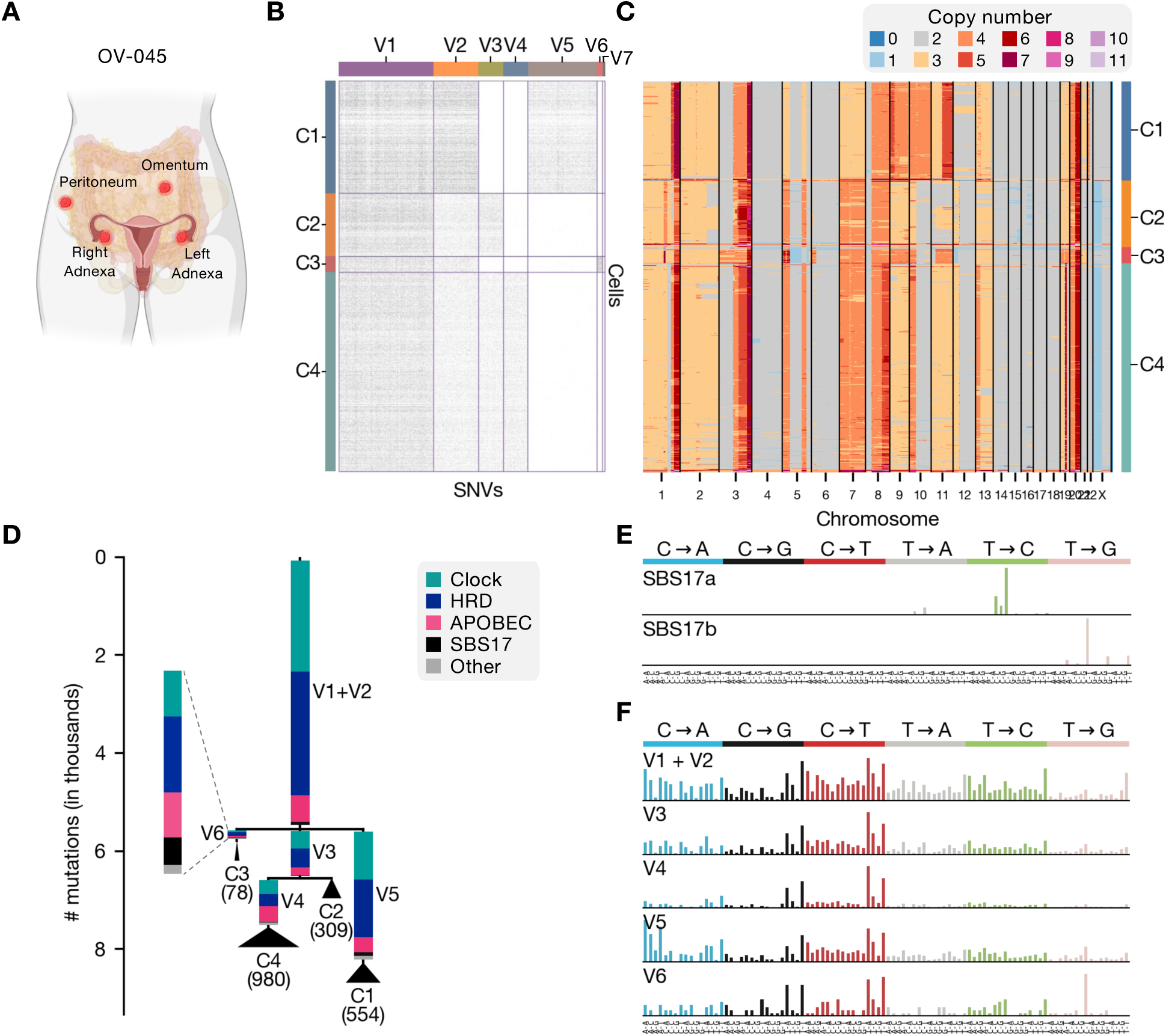
Mutational Signature and Phylogenetic Analysis of HGSOC Sample OV-045. **(A)** Overview of HGSOC OV-045. **(B)** SBMClone clustering results for the ArtiCull filtered SNVs, identifying four cell clusters (C1-C4) and seven SNV clusters (V1-V7). **(C)** Heatmap of the copy number profiles for the cells in OV-045. Each SBMClone-defined cell cluster corresponds to a set of cells with distinct copy-number profiles. **(D)** Phylogenetic tree based on SBMClone SNV clusters. Branch lengths are proportional to the number of mutations. Numbers in parentheses indicate the number of cells in each cluster. **(E)** Reference SBS17a/b mutational signatures. **(F)** Mutation profiles for each SNV cluster/phylogenetic branch.

In addition, we see a clone-specific enrichment of SBS17a/b activity in SNV cluster V6, which is private to cell cluster C3. In V6, SBS17a/b jointly constitute 13% of mutations, while in other SNV clusters, SBS17a is absent and SBS17b is only present in low levels (*<*2%). This shift is observable despite only 78 cells in C3, and 234 SNVs in V6. Interestingly, in a separate longitudinal study of this patient^45^, the clone corresponding to C3 was found to drive recurrence after two subsequent rounds of chemotherapy despite being undetectable in the first recurrence. To confirm the presence of SBS17a/b, we inspected the nucleotide in the −2 genomic position for each SNV (i.e., the 3’ nucleotide immediately preceding the trinucleotide context), which is known to be enriched for A or T in these signatures^1^. The signature decomposition for cluster V6 suggests SBS17a/b accounted for 74% of mutations in the seven trinucleotide contexts associated with SBS17a and SBS17b: C[T→G]{C,G,T} and N[T→G]T, respectively. In these trinucleotide contexts 72% (24*/*33), of the mutations in V6 had an A or T in the −2 position. This represents an enrichment (binomial *p*-value = 0.02) compared to the expected 57% based on the genome-wide A/T proportion^46^. In the other SNV clusters (V1-V5), we also see evidence of an enrichment in the four SBS17a trinucleotide contexts. In these contexts, the signature accounted for 64% of mutations despite low overall exposure values (*<*2%), and 68% (492*/*716) of mutations in these contexts had an A or T in the −2 position (*p <* 0.0001).

## 3 Discussion

We demonstrated how ultra-low-coverage single-cell whole-genome sequencing can reveal the temporal and spatial dynamics of mutational processes during tumor evolution. Mutational signature analysis has been a powerful tool in cancer genomics^1^, but its use has largely been limited to bulk sequencing data. Within single-cell data, signature analysis has mostly focused on copy-number and structural variant signatures^47^. The ability to reliably detect and analyze single-nucleotide variant signatures represents a significant methodological advance. SNV signatures have proven particularly valuable in bulk sequencing studies due to their computational tractability and established connections to both prognostic outcomes and treatment response^4, 5^. By extending these analyses to single-cell data, we enable researchers to study how these clinically-relevant signatures evolve over time and vary across distinct tumor populations.

In context of related work, ArtiCull uses a supervised learning framework similar to past refinement methods (such as DeepSVR^38^) to identify artifactual variant calls based on read and genome features. However, ArtiCull is distinguished by requiring no external training labels. Instead, ArtiCull incorporates evolutionary constraints inferred from copy number alterations (CNAs) to identify a confident subset of artifacts and uses these partial labels to train a supervised classifier. Several previous methods intended for higher-coverage data, such as SCIΦ^31^, Phylovar^48^, SIEVE^49^, and SCIΦN^32^, incorporate evolutionary constraints in single-cell variant calling by simultaneously inferring a phylogeny. ArtiCull by contrast obviates the expense and complexity of simultaneous phylogeny inference.

We demonstrated the broad utility and generalizability of our approach across three distinct cancer types. In PDAC, we inferred a complex phylogeny and tracked changes in mismatch-repair deficiency activity across different lineages during tumor evolution, showing how single-cell approaches can reveal temporal dynamics of mutagenic processes. In cisplatin-treated TNBC PDX models, our method demonstrated the ability to infer expected therapy-induced mutagenesis, validating its capability to capture known effects of treatment. In HGSOC, we showed that ArtiCull removed sample-specific artifacts to enable an analysis of SNV-based clonal structure that was more consistent with copy-number clones.

Our analysis of HGSOC samples uncovered unexpected dynamics of APOBEC3 mutagenesis, a process with established clinical significance in multiple cancer types^50^. APOBEC3 signatures are prominent in breast cancer, where they correlate with poor prognosis and increased metastatic potential^51^, and in cervical cancer, APOBEC3 mutagenesis drives early carcinogenesis and continues to shape tumor evolution throughout progression^52^. However, despite genomic similarities between HGSOC and cancers where APOBEC3 plays a crucial role in progression and treatment response, its significance in HGSOC progression remains largely unexplored. Our observation of both the presence and differing evolutionary dynamics of APOBEC3 mutagenesis in these two patients demonstrates the complex and variable nature of this process in HGSOC. Given the established links between APOBEC3 activity and both treatment response and disease progression in other cancers, these findings warrant deeper investigation into the role of APOBEC3 mutagenesis in HGSOC progression and treatment response.

The detection of clone-specific SBS17 activity demonstrates how single-cell analysis can reveal clone-specific mutational processes that may be masked in bulk sequencing data. SBS17 is rare in HGSOC, detected in only 4% of tumors across major sequencing cohorts^53^. However, our finding that it can emerge late in specific clones suggests that its prevalence may be underestimated in bulk sequencing studies. The subsequent dominance of the SBS17-enriched clone in recurrence highlights the potential importance of understanding how rare or clone-specific mutational processes might influence tumor progression and treatment response. The origin of SBS17 in cancer remains poorly understood. It has been most extensively studied in gastrointestinal cancers, and has been linked in some cases to exposure to 5-fluorouracil chemotherapy and damage from reactive oxygen species^54^. However, whether SBS17a/b reflects a specific mutagenic process relevant to HGSOC progression or treatment response remains an open question.

There remain several methodological limitations and opportunities for future work. Although ArtiCull enables more reliable detection of variants than previous approaches, it still requires multiple supporting reads for variant calling, limiting our ability to study variants unique to individual cells in low-coverage data. Future methodological developments incorporating more sophisticated error models could help overcome this limitation. While we focused on single-nucleotide variants, extending our methods to other mutation types (e.g., dinucleotide variants and small indels) could also provide a more complete picture of ongoing mutagenesis. Additionally, improved methods to quantify shifts in signature exposures along phylogenies with statistical rigor are needed to further refine evolutionary inferences.

The applications of improved single-cell SNV detection extend beyond mutational signature analysis. For example, recent work by McPherson et al.^44^ integrated ArtiCull-filtered variants with copy-number data to reveal complex patterns of whole-genome duplication in HGSOC, using SNVs to provide evidence for multiple independent WGD events within single tumors and infer relative evolutionary timings of these events. This underscores how more accurate SNV detection can deepen our understanding of major genomic events in tumor evolution.

Looking forward, this approach opens new avenues for studying tumor evolution and therapeutic response. Integrating scWGS data with other single-cell modalities (e.g., transcriptomics, epigenomics) could reveal the molecular mechanisms associated with changes in mutational processes. Application to larger cohorts and diverse cancer types may uncover previously unrecognized patterns in how mutational processes shape tumor progression. Ultimately, a better understanding of ongoing mutagenesis could inform strategies to prevent or delay therapeutic resistance and improve long-term outcomes for patients.

## 4 Methods

### 4.1 ArtiCull evolutionary model

In ArtiCull, we define an evolutionary model based on two assumptions that constrain the evolution of single-nucleotide variants (SNVs), copy-number aberrations (CNAs), and their interaction. Here, we describe these assumptions and prove that under this model, non-canonical variants cannot result from any valid phylogeny. Let the *haplotype-specific copy-number profile q* = (**x, y**) of a cell be a pair of vectors where *x_i_* ∈ N represents the maternal copy number for bin *i* and *y_i_* ∈ N the paternal.

#### Assumption 1

*Each distinct haplotype-specific copy-number profile evolves at most once*.

Note that Assumption 1 does not prohibit specific copy-number events occurring more than once, as such homoplasy is likely common and has been documented in multiple cancer types^15, 47, 55–58^. This assumption is implicit in copy-number phylogeny methods based on maximum parsimony and maximum likelihood^59–62^, and has also been used in phylogenetic methods that combine SNVs and CNAs^63–67^.

#### Assumption 2

*A substitution may occur at most once at any position in the genome resulting in an SNV. SNVs may be lost or change multiplicity only due to changes in haplotype-specific copy number*.

The first part of Assumption 2 corresponds to the infinite sites assumption (ISA)^68^ and precludes the same SNV from occurring on different branches of the phylogeny or an SNV from reverting back to the germline state. Due to the short evolutionary time and relatively low mutation rate in cancer development, the ISA has been used commonly to model the evolution of SNVs in cancer^69^. The second part of Assumption 2 reflects that large overlapping copy-number aberrations (deletions and copy-neutral loss-of-heterozygosity events) may result in mutation losses. Assumption (2) as a whole is consistent with previous tumor phylogeny reconstruction methods that combine SNVs and CNAs in both bulk^63, 64, 70^ and single-cell^65–67^ sequencing data. In addition, it underlies SNV-based evolution methods that use the Dollo model^14, 71–74^, which allows SNVs to be gained once but lost multiple times.

A *copy-number clone A* is a set of cells that share an identical haplotype-specific copy-number profile *q_A_*. Hereafter, we refer to copy-number clones as *clones* for simplicity. The *clone-specific cell fraction (CF) c_A_* of an SNV is the proportion of cells in clone *A* that contain a mutation. An SNV is *clonal* with respect to clone A if *c_A_* = 1, subclonal if 0 *< c_A_ <* 1 and absent if *c_A_* = 0. These two assumptions jointly yield the following result.

#### Lemma 1

*For any SNV, there exists at most one clone A in which* 0 *< c_A_ <* 1.

##### Proof

Any copy-number clone *A* defines a subtree *T_A_* of a latent tumor phylogeny *T*, where *T_A_* is the smallest subtree containing all cells in *A*. All extant and ancestral cells in *T_A_* share copy-number profile *q_A_* (Assumption 1), and consequently there are no mutation losses within *T_A_* (Assumption 2). Thus, if an SNV *v* is subclonal with respect to a clone *A*, then *v* was introduced on an edge in *T_A_*. As each mutation is only introduced once (Assumption 2), there can exist at most one clone *A* in which *v* is subclonal.

### 4.2 Identifying non-canonical variants in noisy sequencing data

Due to low per-cell coverage, the observed data provide only noisy estimates of CFs, and thus we cannot trivially identify non-canonical variants. In this section, we describe how we identify non-canonical variants from noisy sequencing data. For a variant in clone *A*, we observe:

- The number of variant reads *v_A_* ∈ ℕ
- The total number of reads *t_A_* ∈ ℕ
- The total copy-number at the locus *n_A_* ∈ ℕ

We model the number of variant reads to be distributed as *v_A_* ∼ Binomial(*p* = *f_A_, n* = *t_A_*), with probability of success *f_A_* over *t_A_* trials. In the absence of sequencing noise, *f_A_* corresponds to the proportion of copies of the locus in the clone containing the variant allele: 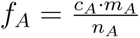 where *m_A_* ∈ [1*, …, n_A_*] is the number of copies of the variant in a cell containing the variant. While the quantity *m_A_* is not directly observed, we may safely make the assumption that *m_A_* = 1 without introducing classification errors (i.e., calling a subclonal variant clonal or a clonal variant subclonal): any subclonal mutations have multiplicity *m_A_* = 1 by Assumptions 1 and 2. Clonal mutations may have a true multiplicity *m_A_ >* 1. However, underestimating *m_A_* would lead to an overestimate of *c_A_*. We thus safely assume *m_A_* = 1.

We introduce a sequencing error frequency of *ɛ*—i.e., in *ɛ* proportion of reads, a variant allele is incorrectly observed as reference allele or a reference allele as a variant. Thus, we observe a variant on a read if either (1) the true base is a variant and we observe it correctly with probability 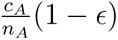; or (2) the true base is the reference base and we incorrectly observe it as a variant with probability 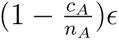. This yields a total

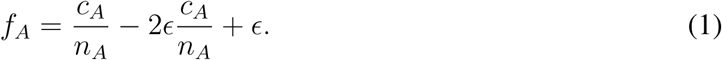

In practice, we use *ɛ* = 0.01. Given this probabilistic model, we define a hypothesis test to identify non-canonical variants between two clones *A* and *B*. Our null hypothesis is that the variant is canonical: the variant is absent in either A or B, or clonal in either A or B. Formally, we have that:

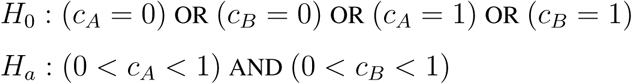

We evaluate this hypothesis test as a disjunction between four tests with null hypotheses 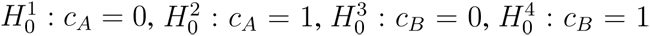. Rejecting all four null hypotheses 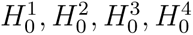 is a rejection of *H*_0_. We compute p-values for each of these hypotheses as:

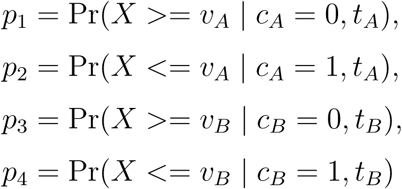

using the Binomial cumulative mass function, with probability of success as given by Equation (1). Tests *p*_1_ and *p*_2_ are independent of tests *p*_3_ and *p*_4_, and thus to achieve an overall false positive rate *α*, we reject each hypothesis if 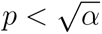. In results, we use *α* = 0.01.

### 4.3 Data processing

All samples were uniformly preprocessed using the DLP+ pipeline available at https://github.com/mondrian-scwgs. SIGNALS^47^ was used for copy-number calling and cell clustering based on copy-number profiles. In bulk datasets (OV-022, OV2295), Mutect2^34^ was used for variant calling. In scWGS, single-cells from were merged together to create a pseudo-bulk genome, then Mutect2 was run on merged data. FilterMutectCalls as part of the GATK pipeline was used to subsequently filter somatic SNVs. As part of benchmarking, Strelka2^36^ and Varscan2^35^ were additionally run on merged scWGS data. SomaticCombiner^37^ was run, using calls from Mutect2, Strelka2 and Varscan2 as input. DeepSVR^38^ was run using calls from Mutect2 as input.

### 4.4 Classifier training and evaluation

124,589 variants from seven DLP+ samples were used for model training, including samples from three triple negative breast cancer (TNBC) tumors (SA501 ^75^, SA535 ^47^, SA1035 ^47^), three high-grade serous ovarian cancer (HGSOC) tumors (SA1047 ^47^, SA1049 ^47^, SA1184 ^47^), and one mammary epithelial cell line (SA609b ^43^). Samples were chosen for training based on visual inspection of pairwise CF distributions (Fig. S4A), selecting cases with at least one pair of sufficiently large clones to allow for reliable identification of non-canonical variants. Copy-number profiles from SIGNALS^45^ were used for computing CFs. Variants were excluded from training if SIGNALS did not return a copy number for the region or if more than 10% of cells in a clone had a copy number different from the dominant/majority copy number for that clone in that region. Variants were labeled as canonical, non-canonical or unlabeled using the hypothesis test with *α* = 0.01, described in Section 4.2, yielding 11,815 non-canonical, 11,386 canonical, and 101,388 unlabeled variants (Fig. S4A-B). For each variant, we extracted fifteen read and genome-level features previously associated with artifactual variants^33^, as defined in Table 1 (Fig. 2A-B, Fig. S4B). For evaluation, traditional cross-validation techniques were not suitable, as the training labels are derived within the algorithm and thus don’t represent a ground truth. Instead, we employed an external validation approach using an independent dataset, OV-022^76^, which provided site-matched bulk sequencing data for HGSOC.

We trained several models, including random forest, logistic regression, linear SVC, gradient-boosted random forest, and multi-layer perceptron, as implemented in scikit-learn, using default parameters. Model performance was evaluated based on AUC relative to the bulk sequencing variant calls from OV-022. The gradient-boosted random forest demonstrated the optimal performance (Fig. S2A). One potential source of errors in identifying non-canonical variants is incorrect assignment of cells to clones. To assess the robustness of ArtiCull to cell-to-clone assignment errors, we introduced varying levels of clone labeling errors (Fig. S2B,C). When label propagation was applied, the model effectively corrected the introduced noise, maintaining stable performance up to a maximum reassignment rate of 16%.

### 4.5 SBMClone and phylogeny construction

To cluster cells and SNVs, SBMClone^39^ was run with settings of 10 restarts and a maximum of 10 SNV clusters (blocks). Phylogenies were constructed based on SBMClone clustering results. Each SNV cluster corresponds to a phylogenetic character, and each cell cluster corresponds to a leaf. A character was considered present if its character density was ≥ 1% in the original mutation matrix, where character density is defined as the proportion of mutation matrix entries corresponding to a given SNV cluster and cell cluster for which the variant allele was observed. Due to the low per-cell coverage (*<* 0.05×), low character densities of 1–5% are expected even when all or most cells in a cluster harbor the SNVs. This procedure yields a binary matrix with cell clusters as rows and SNV clusters as columns. Sets of identical rows were merged, as were sets of identical columns, and rows and columns consisting entirely of 0s were excluded. This procedure resulted in a perfect phylogeny matrix for all analyzed samples^40^, for which a tree was inferred using the perfect phylogeny algorithm. Edge lengths for the phylogenies were assigned according to the number of SNVs in the original clusters.

### 4.6 Mutation signature decomposition

MuSiCal^41^ was used in ‘refitting’ mode for signature decomposition, with the ‘likelihood bidirectional’ method and a threshold of 0.001. Multiple samples exhibited a high frequency of T→A variants occurring within a specific 10-mer context (TTTTTTTT[T→A]AAA). As these variants were detected even in genomically normal cells, they were hypothesized to result from contamination or an artifact introduced during sample preparation. Consequently, they were excluded from further analysis. COSMIC v3.4 signatures^42^ were used for refitting. For each cancer type, available signatures were limited to those previously detected in that cancer type according to the COSMIC database. SBS31 and SBS35 were additionally included for the TNBC PDX samples (Section 2.4) due to known cisplatin exposure. SBS17a/b were included for OV-045 (Section 2.6) based on visual inspection due to their distinct presence in the mutational profiles.

## 5 Data Availability

Single-cell WGS data for SA501 is available in the European Genome-phenome Archive (EGA) under accession number EGAS00001000952. Single-cell WGS and bulk WGS for OV2295 is available under accession number EGAS00001003190. Single-cell WGS for SA609 and SA1035 are available under accession number EGAS00001003190. Single-cell WGS for SA535, SA1049, SA1053, SA1184, and SA906b are available under accession number EGAS00001006343. Single-cell WGS OV-045, OV-046, OV-022, and the PDAC organoids and bulk WGS for OV-022 will be publicly available prior to publication.

## 6 Code Availability

ArtiCull code and trained model are available at https://github.com/shahcompbio/ArtiCull. The pipeline to process DLP+ scWGS is available at https://github.com/mondrian-scwgs. SIGNALS^47^ is available at https://github.com/shahcompbio/signals. Computational analyses were enabled by the Isabl platform^77^.

## Acknowledgments

This work was generously supported by the Nicholls Biondi Chair in Computational Oncology (SPS), a Susan G. Komen Scholar award (GC233085), the Halvorsen Center for Computational Oncology, Cycle for Survival and the Breast Cancer Research Foundation. Additional funding for this work was provided by NCI SPORE (1P50CA247749-01), and NIH CEGS (1RM1HG011014-01).

