## Supplemental Figures and Methods for "Inferring active mutational processes in cancer using single cell sequencing and evolutionary constraints"

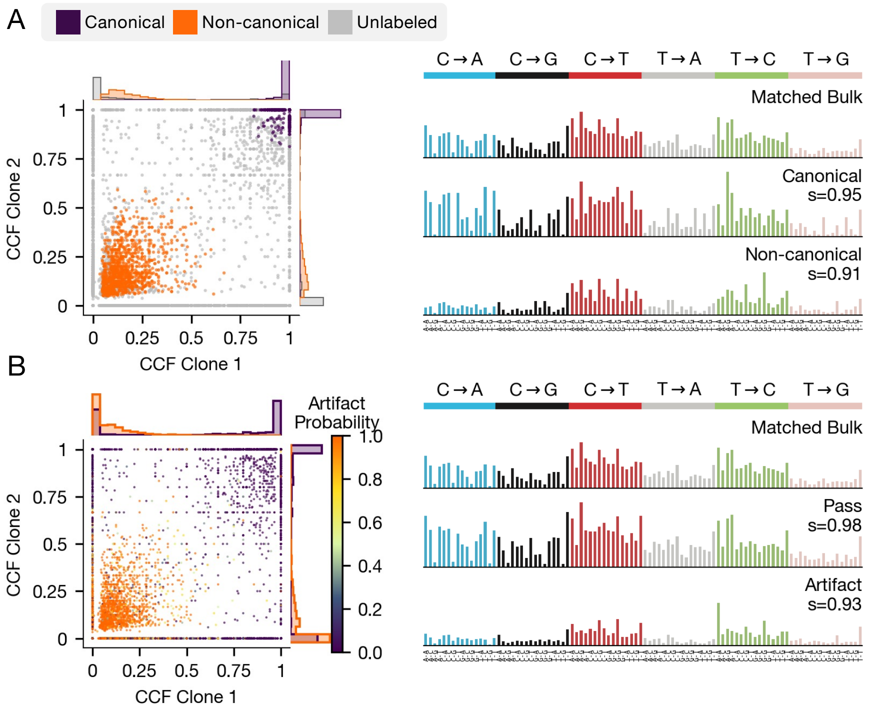


**Figure S1:** **(A)** Pairwise CF comparison for clones 1 and 2 for OV-022. The ArtiCull hypothesis test was used to label the points based on the CCFs. Histograms along the axes show per-clone CF distribution with respect to these labels. (left) Mutation signature distributions for matched bulk, canonical, and non-canonical variants. Cosine similarity ($s$) indicates the similarity between a distribution and the matched bulk distribution. **(B)** The pre-trained ArtiCull model was applied to OV-022. The resulting artifact probabiliities are indicated on the CF distribution by color. Histograms along the axes indicate distributions of mutations identified as artifacts (artifact probability $> 0.5$, in orange) and mutations that pass (prob. $\leq 0.5$ in purple). (left) Mutation signature distributions for bulk, pass, and artifact variants, with cosine similarity as described in panel A.

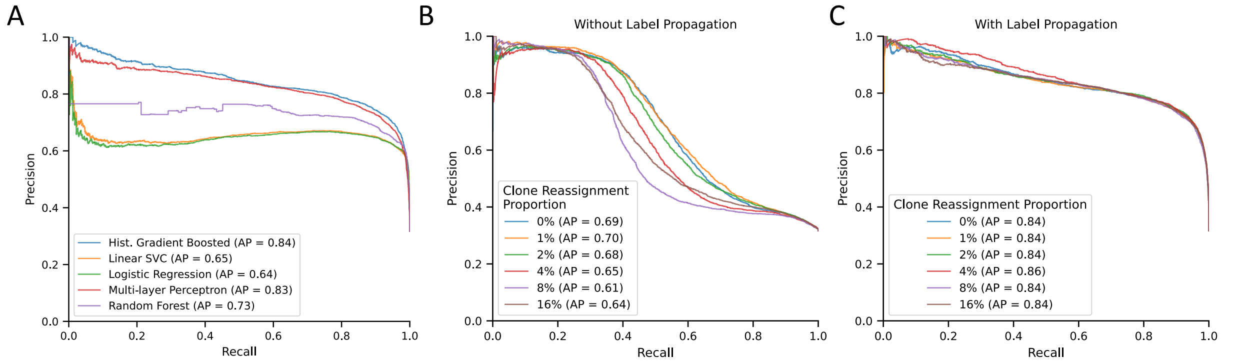


**Figure S2: Additional benchmarking:** (A) Precision-recall curve evaluating different model architectures. (B,C) Precision-recall curve evaluating the effect of reassigning a proportion of cells to different clones to model clone assignment errors. Results were evaluated in a model trained with and without initial label propagation.

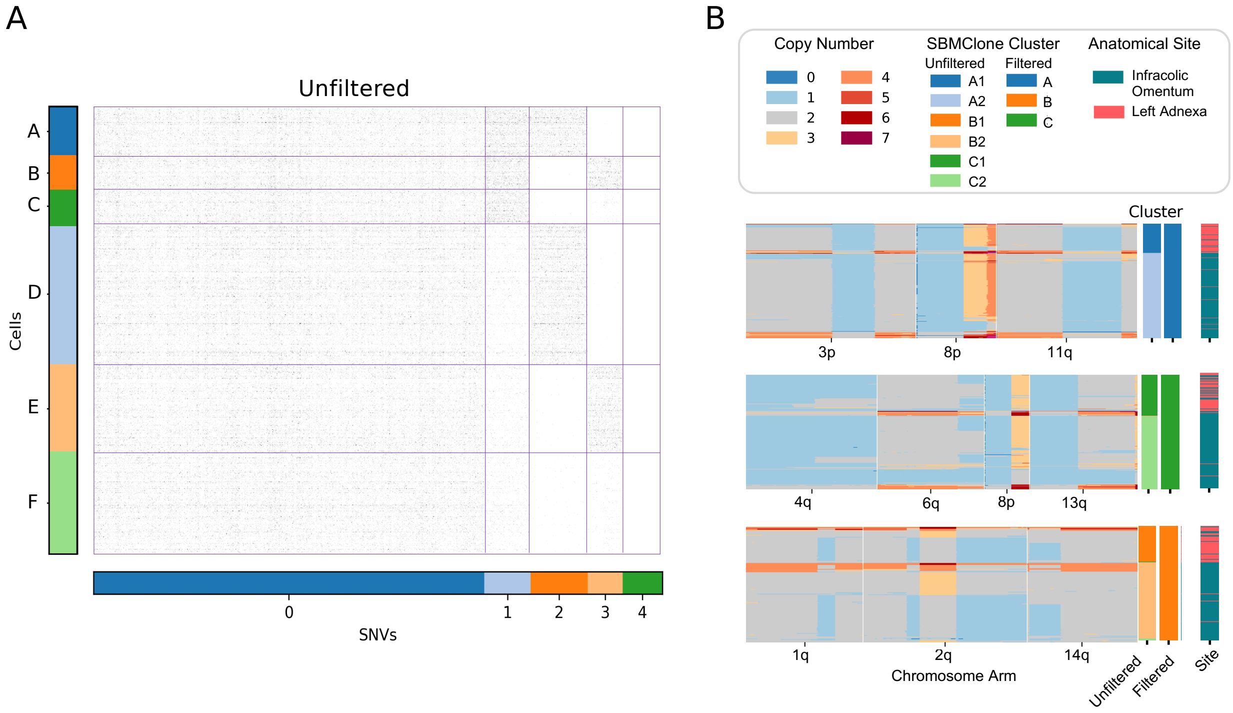


**Figure S3:** (A) SBMClone clustering results for the unfiltered variant call set for OV-046 (Fig. **??**), identifying clusters of cells based on shared SNVs. (B) Copy-number profiles for selected chromosomes in cells, grouped by ArtiCull-filtered SBMClone clusters (Fig. **??**E).

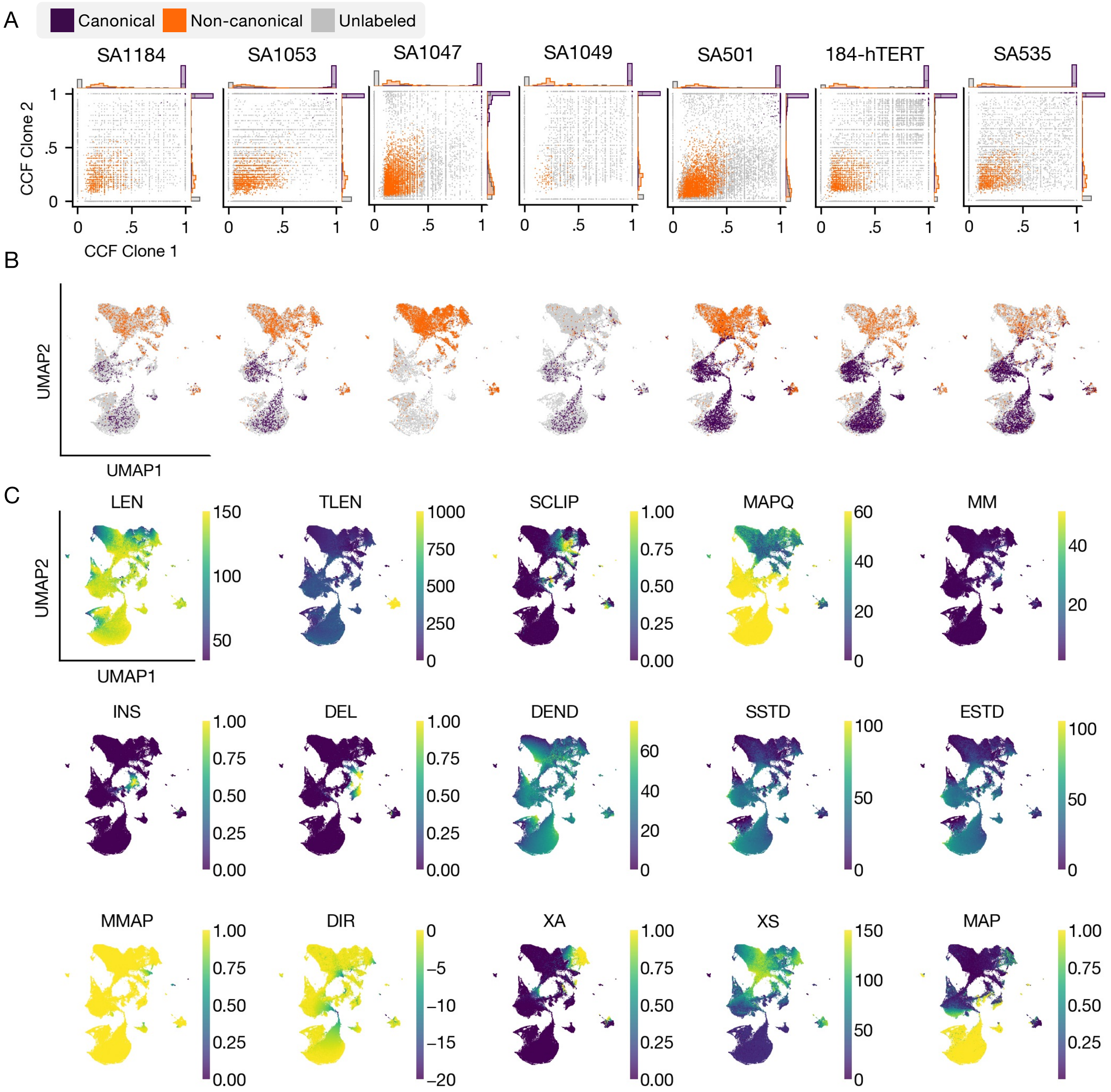


**Figure S4:** (A) Pairwise CF plots for the seven samples used as training data for the model. Each plot visualizes two selected clones per sample, chosen based on clone size (number of cells). Orange points represent Non-Canonical mutations, gray points represent Unlabeled mutations, and purple points represent Canonical mutations. Marginal histograms on the plot axes show the distribution of these mutation labels within each clone. (B) UMAP visualization based on the feature distribution of all training samples. Each plot shows mutations from a given sample, colored by their respective labels (Canonical: purple, Non-Canonical: orange, Unlabeled: gray). Points from other samples are hidden. (C) UMAP visualization of the same feature space as in (B), colored by the value of each of the fifteen model features (yellow: high, purple: low).

#### Supplemental Methods

### S1 Feature Definitions

Let $\mathcal{V}$ be the set of reads $r$ aligning to a locus containing the variant allele. Let $\bar{f}_{\mathcal{V}} = \frac{1}{|\mathcal{V}|} \sum_{r \in \mathcal{V}} f(r)$ correspond to the mean of attribute $f(r)$ over the set of reads $r \in \mathcal{V}$. Let $\sigma^2_{\mathcal{V}}(f)$ be the standard deviation of attribute $f(r)$ over the set of reads $r \in \mathcal{V}$

- LEN $= \bar{L}_{\mathcal{V}}$ where $L(r)$ is the length of read $r$
- TLEN $= \min(\bar{T}_{\mathcal{V}},\ 1000)$ where $T(r)$ is the template length of the read pair containing read $r$.
- MM $= \bar{T}_{\mathcal{V}}$ where $T(r)$ is the template length of the read pair containing $r$
- MAPQ $= \bar{Q}_{\mathcal{V}}$ where $Q(r)$ is the mapping quality of read $r$
- SSTD $= \sigma^2_{\mathcal{V}}(S)$ where $S(r)$ is the start position of read $r$
- EEND $= \sigma^2_{\mathcal{V}}(E))$ where $E(r)$ is the start position of read $r$
- DEND $= \dfrac{1}{|\mathcal{V}|} \displaystyle\sum_{r \in \mathcal{V}} \min\left(p - s(r), e(r) - p\right)$ where $p$ is the position of the variant
- SCLIP $= \bar{C}_{\mathcal{V}}$ where $C(r)$ is an indicator if read $r$ contains soft-clipped bases
- INS $= \bar{I}_{\mathcal{V}}$ where $I(r)$ is an indicator if read $r$ contains inserted bases
- DEL $= \bar{D}_{\mathcal{V}}$ where $D(r)$ is an indicator if read $r$ contains deleted bases
- MMAP $= \bar{M}_{\mathcal{V}}$ where $M(r)$ is an indicator if read $r$ has a properly mapped mate
- XA $= \bar{A}_{\mathcal{V}}$ where $A(r)$ is an indicator if read $r$ has an alternative alignment
- XS $= \bar{G}_{\mathcal{V}}$ where $G(r)$ is the mapping quality of the best alternative alignment
- DIR $= \max(-20,\ \Pr(X \geq f))$ where $f$ is the number of reads with a forward orientation, and $X \sim \text{Binom}(n = |\mathcal{V}|, p = 0.5)$
- MAP $= \dfrac{1}{600} \displaystyle\sum_{\ell=-300}^{300} U(p + \ell)$ where $U(x)$ is the mappability score of position $x$ from UCSC Genome Browser track "wgEncodeCrgMapabilityAlign50mer"
